# High intensity exercise before sleep boosts memory encoding the next morning

**DOI:** 10.1101/2025.01.20.633862

**Authors:** Daniela Ramirez Butavand, Juliane Nagel, Gordon B. Feld, Simon Steib

## Abstract

The importance of sleep for memory consolidation has been extensively studied, but its role for memory encoding remains less well characterized. At the molecular and cellular level, the renormalization of synaptic weights during sleep has received substantial support, which is thought to free capacity to encode new information at the behavioral level. However, at the systems level and behaviorally, support for this process playing a major role for memory function remains scarce. In the current study, we investigated the utility of moderate- and high-intensity evening exercise as a low-cost low-tech intervention to modulate sleep and its influence on subsequent encoding in the morning. Our findings indicate that high-intensity interval training (HIIT) improved post-sleep memory performance with effects lasting up to 24 hours after initial encoding. In addition, we show that especially the early parts of the encoding task were affected by the HIIT intervention, which is in line with increases in synaptic homeostasis being targeted by the exercise. Intriguingly, low-performing participants seemed to benefit more from the HIIT intervention suggesting it not only as a tool for basic research but also as a candidate for applications to boost memory performance in mental disorders or in the elderly. These results provide first evidence that acute exercise can affect learning processes even hours after it occurs.

## Introduction

Sleep plays an important role for efficient memory processing (Feld & Born, 2020; Rasch & Born, 2013; Stickgold & Walker, 2013). During sleep, memories are strengthened and transformed via active systems consolidation, which relies on the repeated reactivation of memory traces that were encoded during prior wakefulness (Diekelmann & Born, 2010). In addition, according to the synaptic homeostasis theory, the brain’s encoding capacity is replenished during sleep (Tononi & Cirelli, 2006, 2014). Taken together, it is suggested that sleep sorts out irrelevant information to enable efficient new learning and preserves relevant information for the long-term (Feld & Born, 2017). Identifying non-invasive interventions that can be used to manipulate the underlying neuronal processes and that enable cost-effective low-tech applications to improve human health and cognition is an important goal of sleep and memory research (Feld & Diekelmann, 2020). At the same time, these simple solutions allow more robust research on the behavioral effects of synaptic homeostasis since they allow larger sample sizes (Button et al., 2013). Utilizing exercise to enhance sleep and the associated cognitive processes is an ideal candidate for such an intervention (Roig et al., 2022).

Behavioral interventions can improve memory, e.g., exposure to stress or novelty in natural settings (Lopes da Cunha et al., 2018; Ramírez Butavand et al., 2020), as well as engaging in physical exercise (Butavand et al., 2023; Jentsch & Wolf, 2020; Roig et al., 2012; van Dongen et al., 2016). Meta-analyses have revealed that acute exercise, defined as a single bout of physical activity, when administered in close temporal proximity to encoding, has a moderate to large effect on episodic and motor memory (Roig et al., 2013; Wanner et al., 2020). Notably, high-intensity interval training (HIIT), i.e., repeated short bouts of high-intensity exercise interspersed with recovery periods at low intensity, that combines high efficacy and low time investment, has been shown to improve memory encoding and consolidation (Kao et al., 2018; Roig-Hierro & Batalla, 2023; R. Thomas et al., 2016).

At the same time, it has been suggested that exercise can modify sleep, although here the evidence is less robust. Nonetheless, meta-analyses concluded that acute exercise affects objective measures of sleep quality and architecture, as well as neurophysiological sleep characteristics (Frimpong et al., 2021; Kredlow et al., 2015; Kubitz et al., 1996; Youngstedt et al., 1997). The most consistent finding is that acute exercise reduces REM sleep and increases non-REM sleep stage 2 (N2) and slow-wave sleep (SWS). Individual studies have also shown that vigorous exercise increases slow wave activity (SWA) and fast spindle activity (Aritake-Okada et al., 2019; Park et al., 2021; Torsvall et al., 1984). Given sleep’s role in memory processing, it is highly plausible that exercise exerts its effects on memory via sleep enhancements. This novel hypothesis is supported by three recent studies (Frimpong et al., 2023; Frisch et al., 2024; Mograss et al., 2020). Although exercise has been shown to enhance sleep-dependent memory consolidation, it has not yet been investigated whether exercise-induced modifications of sleep also enhance memory encoding the following day via the synaptic homeostasis mechanism.

A vast amount of molecular and cellular studies support the synaptic homeostasis theory (Cirelli & Tononi, 2022; Dash et al., 2009; Diering & Huganir, 2018; Hinard et al., 2012; Liu et al., 2010; Schwenk et al., 2014; Suzuki et al., 2020; Vyazovskiy et al., 2008). In humans, the theory is supported by a few studies showing improved memory encoding after sleep versus wakefulness (Mander et al., 2011; Ong et al., 2020) or sleep deprivation (Yoo et al., 2007), accompanied by increased activity in the hippocampus (Ong et al., 2020; Yoo et al., 2007). Moreover, two studies suggested that SWS, and especially the sleep slow oscillation, might be crucial for this effect of sleep in restoring encoding capacity. Antonenko and colleagues (Antonenko et al., 2013) employed transcranial slow oscillation stimulation (tSOS) during a nap, resulting in enhanced SWA and improved encoding of several episodic memory tasks (object recognition, word pairs, and word lists). Conversely, Van Der Werf and colleagues (Van Der Werf et al., 2009) attenuated SWA by acoustic perturbations during nighttime sleep in elderly participants.

This manipulation led to worse post-sleep encoding preceding an image recognition task, accompanied by reduced hippocampal activation during encoding of later recalled items. Both of these interventions require a relatively sophisticated technical setup. Utilizing exercise as a simple intervention to enhance sleep’s restorative effect therefore represents an opportunity to study the basics of synaptic homeostasis theory at the systems level and ultimately develop novel cost-effective interventions targeting this mechanism.

In the present study, we examined the impact of two distinct evening exercise protocols on subsequent nocturnal sleep and memory encoding the following morning. We predicted that both exercise interventions would alter sleep compared to a control condition, resulting in improved memory encoding the next morning.

## Materials and Methods

### Participants

Forty participants (19 female, 21 male, see Table 1 for demographics) completed the three sessions (control, high-intensity interval training [HIIT], and moderate-intensity continuous training [MICT]) of this within-subject design with counterbalanced order (i.e., a total of 120 experimental sessions were conducted). One female participant was subsequently excluded because she did not adhere to our instructions in the control condition (exercised in the morning before the memory task). Demographics were calculated without this participant because she was excluded from all conditions. Data from two HIIT conditions were excluded (1 female, 1 male): one due to technical problems with the online task, and the other because the participant overslept. Data from four MICT sessions were also excluded (3 female, 1 male): two participants overslept, one exercised before the memory task, and one did not finish the memory task. Consequently, the final sample sizes for this within subject-design were 39 for the control condition, 37 for the HIIT condition, and 35 for the MICT condition (the statistical analysis allowed for missing data points, see section *Data Analyses*).

**Table 1:** Demographic variables.

| Characteristics | All (39)<br>Mean (SD) | Female (18)<br>Mean (SD) | Male (21)<br>Mean (SD) |
| --- | --- | --- | --- |
| Age | 25.18 (4.29) | 25.17 (4.58) | 25.19 (4.14) |
| BMI (kg/m<sup>2</sup>) | 22.48 (2.67) | 21.57 (2.68) | 23.26 (2.46) |
| VO<sub>2Max</sub> (ml/min/kg) | 49.44 (10.90) | 43.33 (9.69) | 54.67 (9.15) |
| IPAQ MET | 4618 (2489) | 4845 (2846) | 4424 (2190) |
| PSQI score | 4.64 (1.94) | 4.94 (1.92) | 4.38 (1.96) |

The inclusion criteria required participants to be healthy, physically active (engaging in at least 90 minutes of moderate-intensity exercise per week), non-smoking, native German speakers aged 18-35 years. Additionally, participants needed to have regular sleep-wake cycles (excluding shift workers), non-extreme chronotypes (bedtime between 22:00 and 01:00 and wake-up time between 06:00 and 09:00), and a Body Mass Index (BMI) below 30 kg/m². Participants with medical conditions or using medications or drugs that affect the nervous system or learning ability were excluded. Before starting the study, participants were screened via an online questionnaire, which addressed demographic characteristics, chronotype (Morningness-Eveningness Questionnaire, Adan & Almirall, 1991), subjective sleep quality (Buysse et al., 1989), and general health condition (Goodman et al., 2011).

The study was approved by the local ethics committee of Heidelberg University and all participants gave written informed consent prior to participation in the study. This study was preregistered on the OSF (https://osf.io/ujs8h) and any deviations from the preregistered analyses are indicated below.

### Procedure

Participants visited the lab four times (Figure 1). During the first visit, a graded exercise test (GXT) was performed to measure cardiorespiratory fitness and set exercise intensity levels for the experimental days. At least two days later, participants began the experimental conditions. We used a within-subject design, meaning participants took part in the control, HIIT, and MICT conditions in a counterbalanced order.

**Figure 1:**
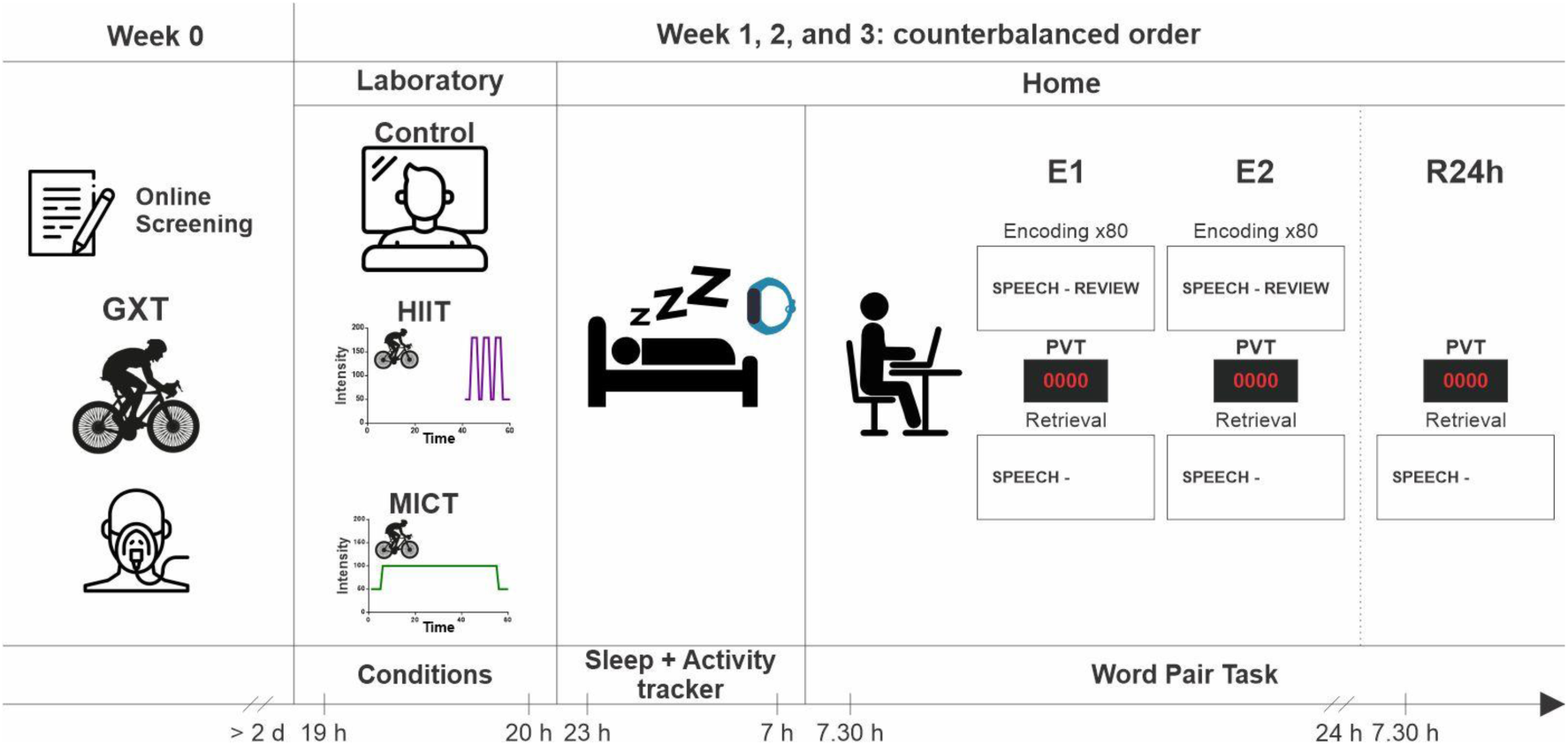
Experimental protocol. Participants first completed the graded exercise test (GXT). At least two days later, they began the first experimental condition (control, HIIT: high-intensity interval training or MICT: moderate-intensity continuous training). After completing the assigned condition, participants went home wearing the activity tracker and slept for eight hours. The following morning, 30 minutes after waking, they performed the memory and vigilance tasks (E1 and E2). Another twenty-four hours later, they completed the final retrieval session (R24h). The protocol was repeated for the two other conditions over the next consecutive weeks, with the order of conditions counterbalanced. The times shown are illustrative, as the protocol was adapted to the participants’ usual bedtime; however, the times between the different steps of the protocol remained the same.

In each condition, participants arrived four hours before their usual bedtime. They performed the assigned condition for one hour, then took a 5-minute shower, put on a wrist-worn actigraphy device, and went home. Participants were instructed to go to bed at their usual time, i.e., approximately three hours after finishing the condition in the laboratory, and to sleep for eight hours.

The next morning, they woke up, ate a provided granola bar, and approximately half an hour after waking, they completed the Stanford Sleepiness Scale (Hoddes et al., 1973) and the St. Mary’s Sleep Questionnaire (Ellis et al., 1981). Immediately afterwards, they performed the memory and vigilance tasks online (see below). After finishing, they continued with their daily routines. The following morning, i.e., approximately 36 hours after the exercise condition, they performed the vigilance task and the retrieval part of the memory task one more time. After completing the last test, actigraphy data collection was stopped.

Participants returned to the laboratory on the same day of the week for the two other conditions. During these weeks, they were instructed to keep their normal physical activity and sleep habits, but were not allowed to exercise, nap, or consume caffeinated beverage after 2 p.m on the experimental days.

### Interventions

*Control*: Participants in the control condition watched a TV documentary for 60 minutes. We selected three episodes from the "Our Green Planet" series by Terra X (in German). The specific episode viewed during each session was also counterbalanced across conditions.

*High-intensity interval training (HIIT)*: In this condition, participants watched a documentary for 40 minutes, and then performed the exercise intervention while continuing to watch the remaining 20 minutes. The exercise was performed on a cycle ergometer and began with a 3-minute warm-up at 50 W. This was followed by three times 3-minute intervals of cycling at 90% of their Wmax, interspersed with two 2-minute intervals at 25% of Wmax. The session concluded with a 4-minute cool-down phase at 50 W.

*Moderate-intensity continuous training (MICT)*: In this condition, participants cycled continuously for 60 minutes at a moderate intensity. The session included a 5- minute warm-up starting at 50 W, with the load progressively increasing to the target intensity. The main 50-minute cycling workout maintained the participants’ heart rate between 60% and 70% of their HRmax, determined during the GXT. Participants watched a TV documentary on a screen in front of them while exercising.

In both exercise conditions, participants were instructed to maintain a pedaling rate of at least 70 rpm throughout the protocol and were allowed to drink water ad libitum. They were also instructed to drink an isotonic gel. Heart rate was continuously recorded, and the rate of perceived exertion (RPE) was assessed with the Borg scale (Borg, 1970).

### Memory and vigilance tasks

*Word Pair Task*: This task was designed to test declarative learning performance using slightly associated word pairs. We created three different lists, each containing 80 word pairs, equivalent in difficulty and used in previous research (Feld et al., 2016). The task was performed online in the participant’s home using JsPsych (De Leeuw, 2015). Each word pair from a list was displayed sequentially on the computer screen for 4 seconds, with a 1-second interval between pairs, followed by a 3-minute version of the psychomotor vigilance task to buffer recency effects (PVT, explained below). After completing the task, the first retrieval session took place, where the cue word (first word in a pair) appeared and the participant had to type in the second target word (encoding performance, E1). This procedure was repeated a second time (E2), with the presentation time for each word pair reduced to 2 seconds. The next morning, participants performed the PVT again, followed by a recall session (24-hour delayed retrieval, R24h).

*Psychomotor Vigilance Task (PVT)*: This task measures participant’s average reaction speed as an indicator of vigilance (Roach et al., 2006). The 3-minute test required participants to press the space bar of their computer as soon as a bright red millisecond timer appeared on the computer screen, starting from 0000 milliseconds. The subject’s reaction time was displayed immediately after pressing the space bar. The mean reaction speed (1 / reaction time in ms) in E1, E2 and R24h was analyzed for each participant in each condition.

### Actigraphy

Sleep data was recorded using activity monitors (GT9X Link, Actigraph Corp, Pensacola, Florida, USA) and analyzed with ActiLife 6 (Actigraph Corp, Pensacola, Florida, USA) software. Sleep times were manually entered based on participants’ reports. From these monitors we obtained latency: time awake after bed onset; efficiency: number of sleep minutes divided by the total number of minutes the subject was in bed; total time in bed (TTB): the total number of minutes in bed; total sleep time (TST): the total number of minutes scored as “asleep”; wake after sleep onset (WASO): the total number of minutes the subject was awake after sleep onset; awakenings: the number of different awakening episodes; average awakening: the average length, in minutes, of all awakening episodes.

### Graded Exercise Test (GXT)

Participants underwent a GXT on a cycle ergometer (Ergoline GmbH, Ergoselect 5, Bitz Germany) with gas exchange measurement (CORTEX Biophysik GmbH, Metalyzer® 3B, Leipzig, Germany) to assess baseline fitness level, including maximal oxygen uptake capacity (VO2max) and maximal power output (Wmax), as well as exercise response parameters (heart rate and blood pressure). The protocol consisted of a 3- minute warm-up at 50 W, followed by stepwise increments of 15 W for women and 20 W for men every minute until subjective exhaustion, and concluded with a 5-minute cool-down phase at 50 W afterwards (Ostadan et al., 2016; Roig et al., 2012). Participants were instructed to keep a pedaling rate of at least 70 rpm throughout the protocol. In addition to the objective measurements, subjective RPE was assessed with the Borg scale.

### Data Analyses

Data were analyzed using R version 4.4.0. Initially, we conducted the pre-registered analyses, i.e., paired t-tests to compare memory performance between the control condition and both the HIIT and MICT conditions at E1. Effect sizes are reported as Cohen’s *d*. We also performed a Pearson’s correlation to assess the relationship between sleep efficiency and memory performance.

To leverage all available data, including incomplete cases, we re-analyzed the data using (generalized) linear mixed models ((G)LMMs), which offer increased statistical power and provide a more robust analysis of repeated measures. Generally speaking, (G)LMMs allow to model random intercepts and random slopes (i.e., intercepts and slopes modelled individually for, e.g., each participant or item). Traditional analyses ignore these features of the data and, thus, (G)LMMs provide more accurate estimates of the variance components contained in the data (Barr et al., 2013; Nebe et al., 2023). These (G)LMMs were also used for the exploratory analyses. We implemented (G)LMMs using the lme4 package in R (version 1.1.35, Bates, 2014), and *p*-values were computed using the lmerTest package (version 3.1.3, Kuznetsova et al., 2017), applying Satterthwaite’s degrees of freedom method. For LMMs (only for the sleep and PVT data, where data was not binary), reported results include *t*-values, degrees of freedom, and *p*-values. GLMMs were used to evaluate the memory data at the trial level (i.e., unaggregated), since these were binary (correct vs. incorrect). Analyzing these data without previous aggregation allowed us to include random effects of the items and thus account for variance in the data stemming from idiosyncratic features of the individual word pairs. For GLMMs, reported results include *z*-values and *β*-weights in log-odds. All predictors were dummy coded: conditions (reference level: control), test sessions (reference level: E1), encoding time (reference level: early), and performance levels (reference level: low-performers). In our model equations, (…|subject) and (…|item) denote random effects by participants and individual word pairs, respectively. For each analysis, we first tried to fit a maximal model (Barr et al., 2013). In cases where the model failed to converge, we addressed the issue by initially simplifying the random effects structure, specifically by removing interaction terms in the random slopes. For models resulting in singular fits, we further examined the correlation estimates among random effects. Random effects terms with correlations approaching ±1 were identified and removed to achieve a well-specified and interpretable model. Once the best-fitting model was identified, we conducted model comparisons to determine whether a simpler model without the interaction term between the fixed effects provided a fit comparable to the more complex model that included the interaction. These comparisons were performed using the ANOVA function, relying on the likelihood ratio test (LRT) and following the recommendation of Matuschek et al. (Matuschek et al., 2017), we used a significance threshold of αLRT = 0.2.

To further explore the effects of experimental conditions and their interactions, as well as to obtain pairwise comparisons, we used estimated marginal means using the emmeans package in R (version 1.10.5, Lenth, 2024).

## Results

### Memory Results

In our preregistered analysis approach, the control and HIIT conditions did not differ regarding the amount of word pairs recalled during the first morning learning session (t-test comparing encoding performance at E1, *t(36)* = 1.50, *p* = 0.143, Cohen’s *d* = 0.25, 95% CI [-0.08, 0.57]). Similarly, the control and MICT conditions at E1 also did not differ regarding the amount of correct word pairs (*t(34)* = 0.23, *p* = 0.822, Cohen’s *d* = 0.04, 95% CI [-0.29, 0.37]).

As indicated in the Data Analyses section we also used more powerful statistical methods and conducted a generalized linear mixed model that included binary data for each item during all measurement points (predictor *test*: E1, E2, and R24h) and for all participants (including those with missing data points as detailed in the methods section). The best-fitting GLMM for this analysis with a logit link function was: *memory performance (0 or 1) ∼ condition + test + (1 + condition | subject)* + *(1 + condition | item)*.

The results indicated overall that participants in the HIIT condition recalled more word pairs than in the control condition (*β* = 0.27, *SE* = 0.13, *z* = 2.02, *p* = 0.043, see Fig. 2). In contrast, participants in the MICT condition did not recall significantly more (or less) word pairs than in the control condition (*β* = 0.07, *SE* = 0.15, *z* = 0.46, *p* = 0.647). In test sessions E2 and R24h, participants recalled significantly more word pairs than in test session E1 (E2: *β* = 1.50, *SE* = 0.04, *z* = 38.11, *p* < 0.001; R24h: *β* = 1.10, *SE* = 0.04, *z* = 28.05, *p* < 0.001).

**Figure 2:**
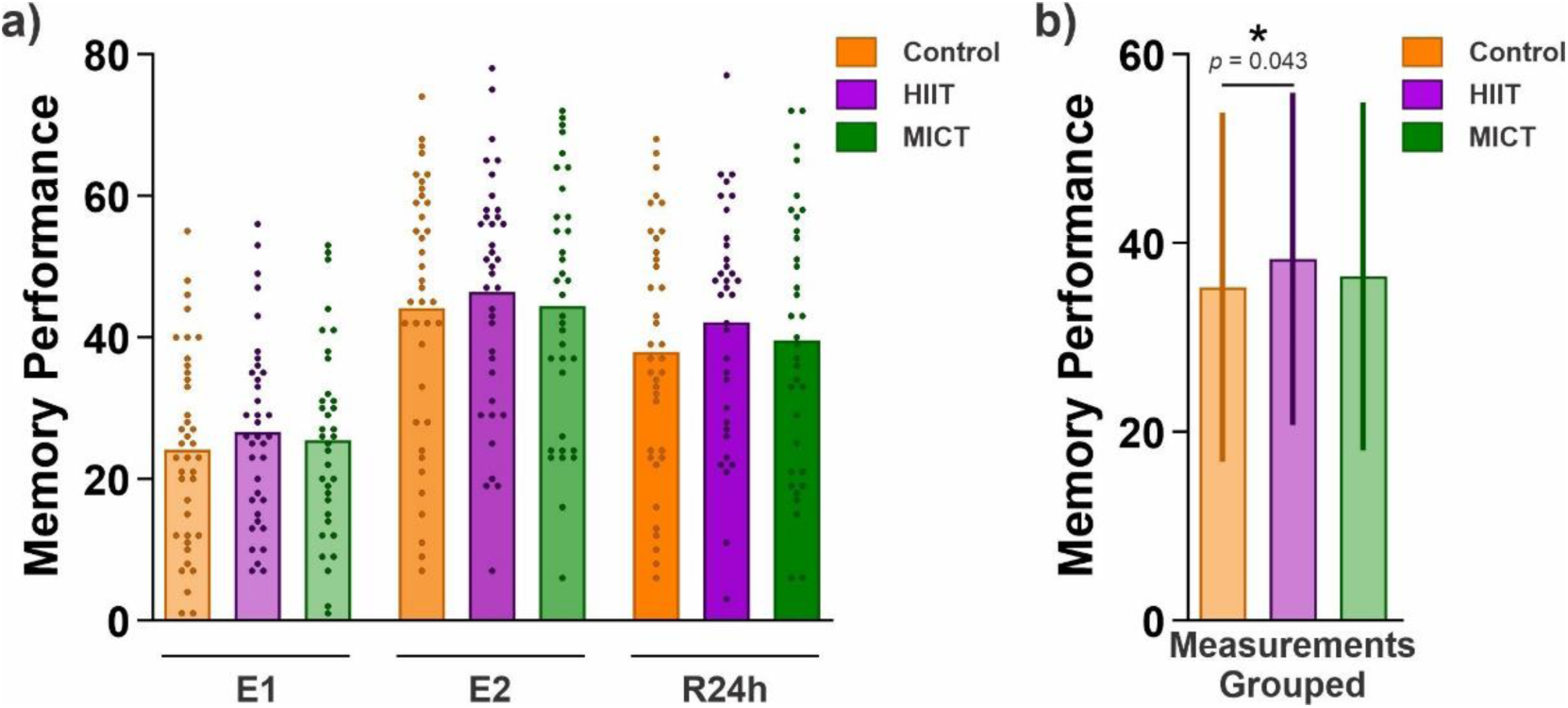
Positive effect of HIIT on memory performance. **a)** Memory performance (i.e., the number of correctly recalled word pairs in each test at E1, E2 and R24h), is shown as the mean for each condition (Control, HIIT, and MICT) plus the performance of individual participants represented by dots. **b)** The measurements across testing time points were averaged for each condition. The performance is shown as the mean ± SD, note that the effect of interest is within-subject. The generalized linear mixed model revealed that participants in the HIIT condition recalled more word pairs than in the control condition. Asterisks represent significance at α = 0.05.

### Objective vigilance

Regarding our test of objective vigilance, we found no significant differences in reaction speed between the conditions in the PVT; however, participants were significantly slower in E2 compared to E1 across all conditions. Detailed statistics are provided in the supplementary information (Supplementary Table 1).

### Sleep Results

We evaluated the effect of exercise on sleep parameters measured by actigraphy. The best-fitting LMM used for analyzing each sleep variable was specified as: *sleep variable ∼ condition + (1 | subject)*. The results revealed that participants spent approximately 8 hours in bed as instructed, with no significant differences in time spent in bed between the control condition and both the HIIT and MICT conditions (see Table 2, TTB). In addition, they spent approximately 6.5 hours sleeping according to sleep latency and awakenings recorded by the actigraphy device, with no significant differences between conditions (see Table 2, TST, WASO, # awakenings, and Avg.awakenings). No significant differences were found in any other sleep parameters between the control condition and the HIIT and MICT conditions. Results for the second night’s sleep (before R24h) are provided in the supplementary materials (Supplementary Table 2).

**Table 2:**
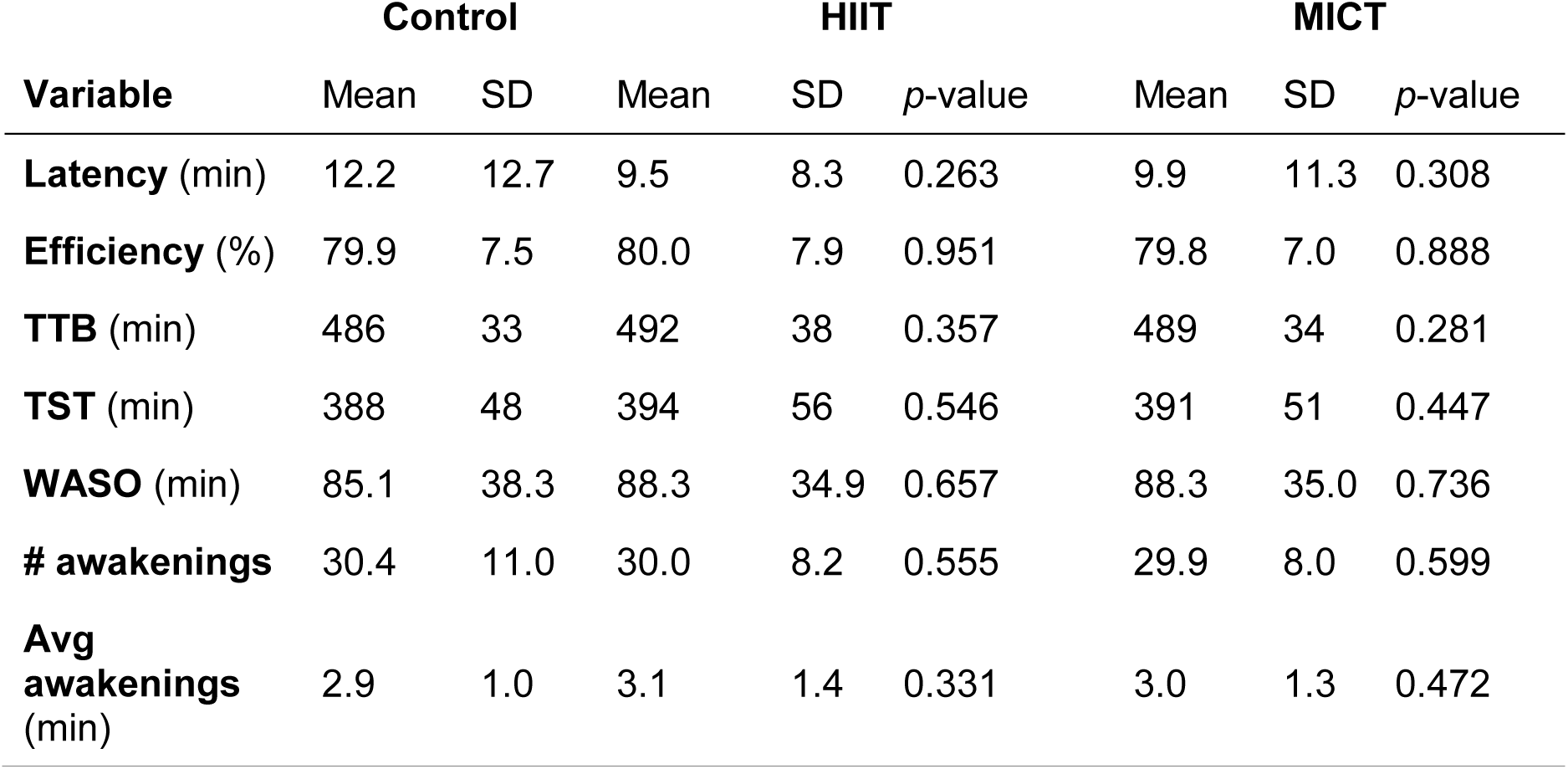
Sleep Variables across experimental conditions. Means and standard deviations (SD) for sleep variables (Latency, Efficiency, Total Time in Bed [TTB], Total Sleep Time [TST], Wake After Sleep Onset [WASO], number [#] of awakenings, and average awakenings) are shown for each condition. In addition, the *p*-values from the comparisons between conditions are displayed.

We also conducted the pre-registered correlation analysis between sleep efficiency and the amount of word pairs recalled at time point E1. There was no significant correlation between these two variables (r = -0.002, *p* = 0.981; see Supplementary Fig. 1).

### Exploratory Results

We conducted two exploratory analyses. First, we investigated whether exercise specifically influenced the initial phase of memory encoding, as would be expected according to the synaptic homeostasis hypothesis. To assess this, we divided the encoding phase into two halves: the first 40 word pairs (early encoding) and the second 40 word pairs (late encoding), analyzing memory performance for each encoding time separately (predictor *encoding time*: early and late). The best-fitting GLMM for this analysis with a logit link function was: *memory performance (0 or 1) ∼ condition * encoding time + (1 + condition + encoding time | subject)* + *(1 | item)*. We first analyzed E1, as adding E2 and R24h to the analysis would make it hard to interpret due to the added interaction. Participants recalled significantly more word pairs from the first half of word pairs encoded during the E1 test in the HIIT condition than in the control condition (*β = 0.46, SE = 0.14, z = 3.35, p < 0.001*, see Fig. 3a), which was not the case for the second half (condition x encoding time interaction: *β = -0.39, SE = 0.13, z = -3.08, p = 0.002*). No significant difference in the amount of word pairs correctly recalled was found for the first half between the control and MICT conditions (*β = 0.26, SE = 0.17, z = 1.57, p = 0.117;* condition x encoding time interaction: *β = -0.42, SE = 0.13, z = -3.20, p = 0.001*). In addition, word pairs retrieved by participants in the control condition did not differ significantly for the two halves (*β = 0.15, SE = 0.11, z = 1.38, p = 0.167*). Results of additional pairwise comparisons revealed no other significant effects. These results are included in the supplementary materials (Supplementary Table 3).

**Figure 3:**
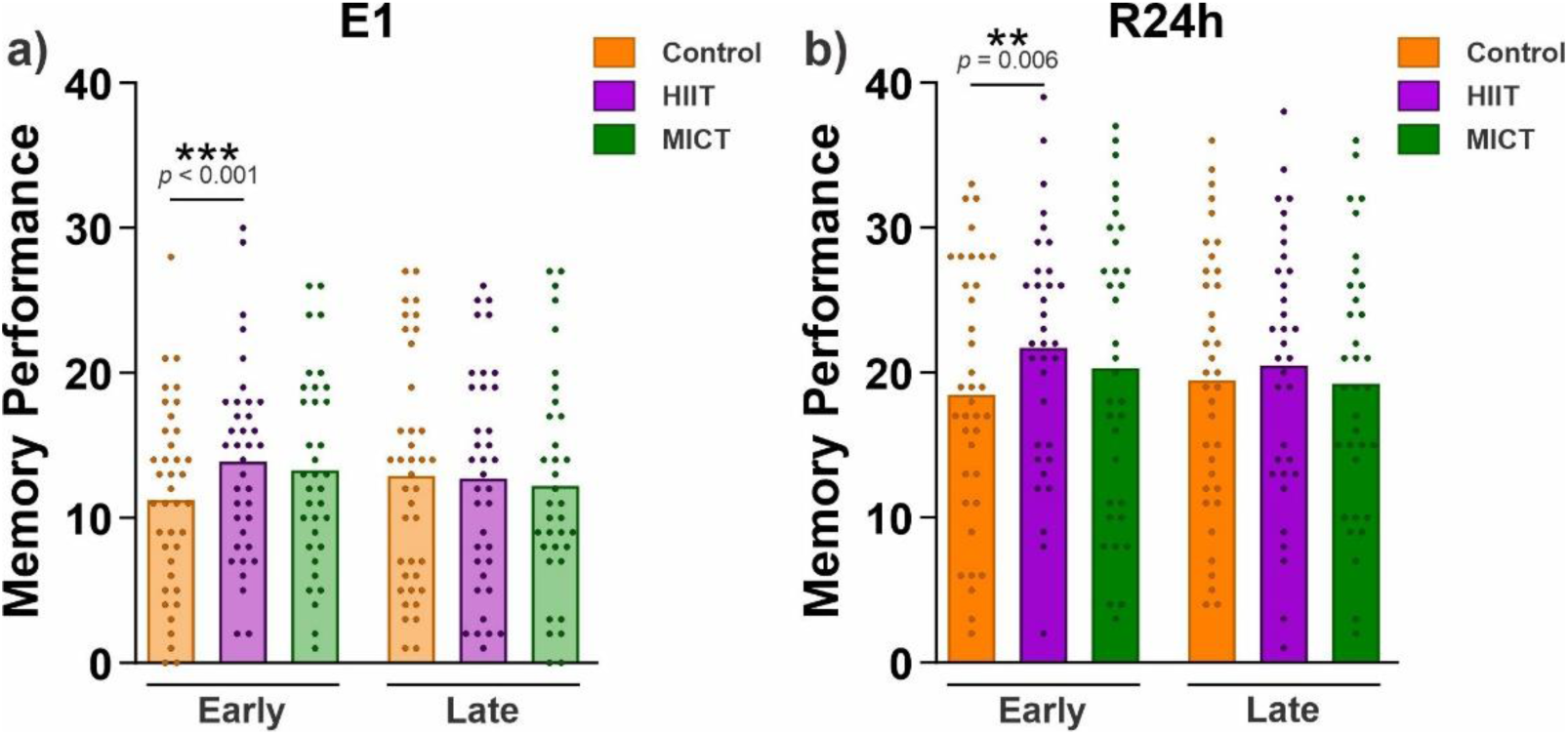
Participants’ memory performance across conditions, separated into first and second halves of encoding. Memory performance for the first (early) and second (late) halves of words encoded in E1 during the **(a)** E1 and **(b)** R24h tests. Data are shown as the mean for each condition and each dot represents the performance of individual participants. The generalized linear mixed models revealed that participants recalled more word pairs from the first half in the HIIT condition than in the control condition in both tests: E1 and R24h. The results for E2 are provided in Supplementary Table 4. Asterisks represent significance at α = 0.05.

Next, we analyzed memory performance in the R24h test for the same words encoded in the first and second halves of the initial encoding session (see Fig. 3b). The pattern of results of this analysis was similar to that of the analysis in E1. Participants in the R24h test recalled significantly more word pairs only from the first half encoded in the HIIT condition compared to the control condition (*β* = 0.43, *SE* = 0.16, *z* = 2.77, *p* = 0.006; condition x encoding time interaction: *β* = -0.30, *SE* = 0.13, *z* = -2.26, *p* = 0.024). There was also no significant difference in the number of word pairs correctly recalled from the first half between the control and MICT conditions (*β = 0.22, SE = 0.18, z = 1.19, p = 0.234;* condition x encoding time interaction: *β = -0.28, SE = 0.13, z = -2.09, p = 0.037*). A similar pattern of results was found for E2 (see Supplementary Table 4).

In the second exploratory analysis, we tested whether exercise had different effects on low- and high-performing participants. To examine this, we divided participants into low- and high-performing groups based on a post hoc median split of their performance in the control condition during E1 (predictor *performer*: low and high). The best-fitting GLMM for this analysis with a logit link function was: *memory performance (0 or 1) ∼ condition * performer + (1 + condition | subject)* + *(1 | item)*. The results revealed that low-performing participants recalled significantly more word pairs in the HIIT condition than in the control condition (*β* = 0.52, *SE* = 0.16, *z* = 3.15, *p* = 0.002, see Fig. 4a), which was not the case for high-performing participants (condition x performer interaction: *β* = -0.51, *SE* = 0.23, *z* = -2.23, *p* = 0.026). Low-performing participants did not differ in the number of word pairs recalled in the control and MICT conditions (*β* = 0.25, *SE* = 0.22, *z* = 1.15, *p* = 0.249; condition x performer interaction: *β* = -0.37, *SE* = 0.29, *z* = -1.26, *p* = 0.209). Additionally, high-performing participants recalled significantly more word pairs than low-performing participants (*β* = 1.71, *SE* = 0.22, *z* = 7.78, *p* < 0.001). Results of additional pairwise comparisons revealed no other significant effects. These results are included in the supplementary materials (Supplementary Table 5). A similar pattern of results was found in E2 (Supplementary Table 6) and in R24h (see Fig. 4b and Supplementary Table 7).

**Figure 4:**
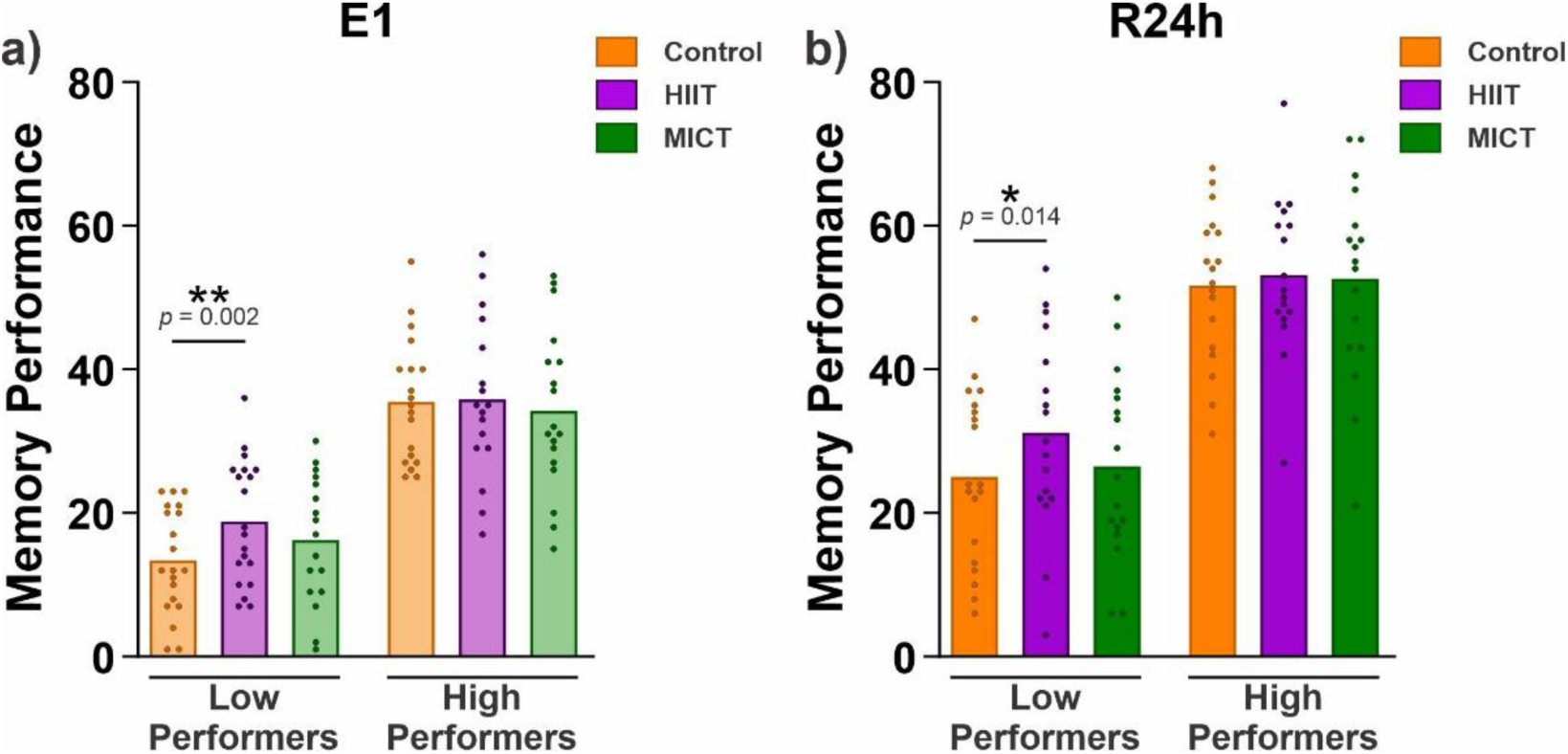
Participants’ memory performance across conditions, separated into low- and high-performers. Memory performance for low- and high-performing participants in the **(a)** E1 and **(b)** R24h tests. Data are shown as the mean for each condition and each dot represents the performance of individual participants. The generalized linear mixed models revealed that low-performing participants recalled more word pairs in the HIIT condition than in the control condition in E1 and R24h. Asterisks represent significance at α = 0.05. The statistical results for E2 are shown in the Supplementary Table 6 and the full report for R24h in the Supplementary Table 7.

## Discussion

In this study, we investigated the effects of two evening exercise interventions on post-sleep memory encoding of word pairs and demonstrated the utility of high-intensity exercise for basic systems level research on synaptic homeostasis and for potential future applications. To our knowledge, this is the first study of this kind. To achieve a larger sample size than has usually been reported in the fields of exercise and sleep (Antonenko et al., 2013; Aritake-Okada et al., 2019; Dworak et al., 2008; Van Der Werf et al., 2009), we used a novel approach that combines online (De Leeuw, 2015) and in-lab experimental methodologies. Consistent with our preregistered hypothesis, our findings indicate that an acute bout of intense exercise (HIIT) three hours before bedtime significantly enhances memory performance the following morning. However, this effect only emerged when we employed more robust statistical methods (not preregistered), suggesting a potential limitation in the sensitivity of the original analytical approach. Contrary to our predictions, this improvement was not observed following an acute bout of moderate exercise (MICT). Actigraphy-derived sleep data did not show any significant changes to sleep due to the interventions. Exploratory analyses revealed that participants in the HIIT condition encoded the early items of the word pair task significantly better than in the control condition. The early items were also better recalled after 24 hours. Furthermore, low-performing participants exhibited improved memory encoding in the HIIT condition compared to the control condition, which extended to enhanced memory retention after 24 hours. This effect was not present in high-performing participants, nor did we find evidence for such an effect in the MICT condition.

Previous studies have investigated how exercise administered immediately before memory encoding can impact memory performance (Frith et al., 2017; Loprinzi et al., 2019). However, these studies did not schedule sleep in-between to observe its impact on post-sleep memory encoding. According to the sleep and synaptic homeostasis hypothesis (Tononi & Cirelli, 2003, 2006), sleep architecture changes, particularly in NREM sleep properties such as SWA and spindles, play a crucial role in restoring encoding capacity (Antonenko et al., 2013; Ong et al., 2020; Van Der Werf et al., 2009; Yoo et al., 2007). A plausible explanation for our findings is that HIIT may enhance encoding by modulating sleep architecture, potentially through increased slow waves and/or spindles. Slow waves (activity in the 1-4 Hz range) facilitate synaptic downscaling and restore encoding capacity, while spindles support hippocampal-to-neocortical transfer of reactivated information, freeing hippocampal resources for subsequent learning (Feld & Born, 2017; Rasch & Born, 2013; Tononi & Cirelli, 2003, 2006). Their putative enhancement by HIIT may be driven by exercise-induced elevation in core body temperature, which leads to a greater distal-to-proximal skin temperature gradient during sleep, linked to increases in both spindle activity and SWA (Aritake-Okada et al., 2019). Supporting these hypotheses, evidence suggests that high-intensity exercise is effective in increasing N2 and SWS, stages during which spindles and slow waves predominantly occur (Aritake-Okada et al., 2019; Frisch et al., 2024; Park et al., 2021; Torsvall et al., 1984).

Although we expected with both exercise interventions, only high-intensity exercise significantly improved memory performance, while moderate exercise did not. While some studies have reported increased SWA and SWS following moderate exercise (Hayashi et al., 2014, n = 9; Park et al., 2021, n = 9; Yoshida et al., 1998, n = 5), these findings were based on small sample sizes, and in some cases, the increases were not statistically significant. A potential explanation for our findings is that MICT may not have induced meaningful changes in sleep architecture sufficient to enhance memory encoding the following morning. In particular, a single bout of moderate-intensity exercise may not have elevated core body temperature enough in our fit participants to affect SWA and SWS. Interestingly, Aritake-Okada and colleagues’ study (Aritake-Okada et al., 2019) found significant increases in SWS following four sessions of moderate-intensity exercise in a sedentary population, highlighting the role of exercise volume and participant fitness. In addition, some studies have found that SWS elevation occurs only after high-intensity, but not moderate-intensity, exercise (Dworak et al., 2008). Further well-powered studies with polysomnography are needed to better understand how different exercise intensities affect sleep architecture and their subsequent impact on memory.

In studies measuring sleep with both actigraphy and polysomnography, exercise-induced changes, such as reduced REM sleep and increased non-REM sleep, were found only with polysomnography (Aloulou et al., 2020, n = 12; Myllymäki et al., 2011, n = 11; C. Thomas et al., 2020, n = 8). Although our study was run using a much larger sample (at least 35 subjects per condition, within-subjects) allowing to detect behavioral results with more precision this led to resource constraints and only allowed us to collect actigraphy and not polysomnographic data. Hence, it may be unsurprising that we did not find significant differences in any of the sleep variables measured with actigraphy. While this provides robust evidence that actigraphy lacks sensitivity to detect exercise-driven changes in sleep it does not necessarily indicate that HIIT has no measurable effect on sleep architecture, since REM and NREM sleep can only be reliably differentiated with polysomnography and this more involved technique allows identifying slow waves and sleep spindles. Notably, we did not observe any sleep disturbances resulting from the evening exercise interventions since sleep quality remained high, which is important to consider, when considering such interventions for broader use. In sum, polysomnography is needed to determine effects of exercise on sleep.

To gain further insight into the influence of exercise on morning encoding, we conducted two exploratory analyses. We found better early encoding in the HIIT group than in the control group. Notably, the enhanced early encoding of the HIIT group led to better memory performance of those early-encoded pairs also after 24 hours. A possible explanation is that the second half of the word pairs retroactively interfered with the memory of the first half, a phenomenon observed in previous studies (Lechner et al., 1999; Robertson, 2012). The HIIT intervention may counteract this interference by enhancing synaptic downscaling leading to more robust encoding of the early items and stronger traces as indicated by better memory performance. Second, we found that low-performing participants showed improved memory encoding in the HIIT condition compared to the control condition, which also led to better memory performance in the 24-hour delay test. Previous studies have examined the effects of sleep on memory consolidation in low- and high-performers, yielding mixed results: Diekelmann and colleagues (Diekelmann et al., 2010) found a positive effect of sleep only for low memory performers, while Tucker and Fishbein (Tucker & Fishbein, 2008) observed a positive effect only for high-performers. One study has shown that a closed-loop tACS intervention, designed to enhance slow-wave oscillations, can prevent retroactive interference compared to sham stimulation, particularly in poor encoders (Jones et al., 2023). Although the protocol differs from ours, the HIIT intervention may similarly help low-encoders compensate poor encoding abilities. While these findings should be interpreted with caution, they are promising, as this simple intervention, potentially feasible for home use, could offer a practical way to help individuals with memory impairments improve their cognitive function.

Although our pre-registered analyses were not significant, the more powerful GLMMs, which utilized the full variance of the repeated measurements, proved to be sensitive in detecting differences. Improving the statistical analysis can be a reason to deviate from a preregistration (Lakens, 2024). Importantly, we preregistered our study to promote transparency and replicability, even though our investigation explored a novel hypothesis; we therefore place a decent amount of trust in our results that nonetheless need to be replicated independently in the future. We did not observe any effects on sleep parameters; therefore, our attribution of the behavioral effects on memory to the interaction between exercise and sleep remains a speculation until supported by polysomnography. Nonetheless, our innovative approach of combining in-lab and online experiments allowed us to include a larger sample size than usual, and compared two types of exercise interventions, optimizing the informative value of our data within the constraints of available resources (Lakens, 2022).

In conclusion, we demonstrate a positive effect of an acute evening bout of HIIT on post-sleep memory performance, especially for low-performing individuals. This is the first study to explore this specific interaction, laying important groundwork for future research to further investigate the mechanisms underlying these findings. Notably, this approach offers a simple way to modify sleep and assess its impact on brain plasticity and memory encoding, which may offer new insights into the mechanisms of the synaptic renormalization. Moreover, these findings may establish the basis for practical interventions readily applicable in populations suffering from both sleep and memory disorders.

## Acknowledgements

This research was supported by the Health + Life Science Alliance Heidelberg Mannheim and received state funds approved by the State Parliament of Baden-Württemberg. This research was also supported by an Emmy-Noether-Grant (FE 1617/2-1) to Gordon B. Feld.

## Author Contributions

DRB: developed the study concept and study design, generated the study material, performed data collection, performed the data analysis and interpretation, drafted and revised the manuscript. JN: supported the generation of the study material and the data analysis and interpretation. GBF: developed the study concept and design, supervised data analysis and interpretation, revised the manuscript. SS: developed the study concept and design, supervised data analysis and interpretation, revised the manuscript. All authors approved the final version of the manuscript for submission.

## References

1. Adan, A., & Almirall, H. (1991). Horne & Östberg morningness-eveningness questionnaire: A reduced scale. Personality and Individual Differences, 12(3), 241–253. 10.1016/0191-8869(91)90110-W

2. Aloulou, A., Duforez, F., Bieuzen, F., & Nedelec, M. (2020). The effect of night-time exercise on sleep architecture among well-trained male endurance runners. Journal of Sleep Research, 29(6), e12964. 10.1111/jsr.12964

3. Antonenko, D., Diekelmann, S., Olsen, C., Born, J., & Mölle, M. (2013). Napping to renew learning capacity: Enhanced encoding after stimulation of sleep slow oscillations. European Journal of Neuroscience, 37(7), 1142–1151. 10.1111/ejn.12118

4. Aritake-Okada, S., Tanabe, K., Mochizuki, Y., Ochiai, R., Hibi, M., Kozuma, K., Katsuragi, Y., Ganeko, M., Takeda, N., & Uchida, S. (2019). Diurnal repeated exercise promotes slow-wave activity and fast-sigma power during sleep with increase in body temperature: A human crossover trial. Journal of Applied Physiology, 127(1), 168–177. 10.1152/japplphysiol.00765.2018

5. Barr, D. J., Levy, R., Scheepers, C., & Tily, H. J. (2013). Random effects structure for confirmatory hypothesis testing: Keep it maximal. Journal of Memory and Language, 68(3), 255–278.

6. Bates, D. (2014). Fitting linear mixed-effects models using lme4. arXiv Preprint arXiv:1406.5823. https://discourse.mc-stan.org/uploads/short-url/lAx43nMH1ZjNqyRjZzLveIXUDAR.pdf

7. Borg, G. (1970). Perceived exertion as an indicator of somatic stress. Scandinavian Journal of Rehabilitation Medicine. https://psycnet.apa.org/record/2018-29834-001

8. Butavand, D. R., Rodriguez, M. F., Cifuentes, M. V., Miranda, M., Bauza, C. G., Bekinschtein, P., & Ballarini, F. (2023). Acute and chronic physical activity improves spatial memory in an immersive virtual reality task. Iscience, 26(3). https://www.cell.com/iscience/fulltext/S2589-0042(23)00253-5

9. Button, K. S., Ioannidis, J. P. A., Mokrysz, C., Nosek, B. A., Flint, J., Robinson, E. S. J., & Munafò, M. R. (2013). Power failure: Why small sample size undermines the reliability of neuroscience. Nature Reviews Neuroscience, 14(5), 365–376. 10.1038/nrn3475

10. Buysse, D. J., Reynolds III, C. F., Monk, T. H., Berman, S. R., & Kupfer, D. J. (1989). The Pittsburgh Sleep Quality Index: A new instrument for psychiatric practice and research. Psychiatry Research, 28(2), 193–213.

11. Cirelli, C., & Tononi, G. (2022). The why and how of sleep-dependent synaptic down-selection. Seminars in Cell & Developmental Biology, 125, 91–100. 10.1016/j.semcdb.2021.02.007

12. Dash, M. B., Douglas, C. L., Vyazovskiy, V. V., Cirelli, C., & Tononi, G. (2009). Long-term homeostasis of extracellular glutamate in the rat cerebral cortex across sleep and waking states. Journal of Neuroscience, 29(3), 620–629.

13. De Leeuw, J. R. (2015). jsPsych: A JavaScript library for creating behavioral experiments in a Web browser. Behavior Research Methods, 47, 1–12.

14. Diekelmann, S., & Born, J. (2010). The memory function of sleep. Nature Reviews Neuroscience, 11(2), 114–126. 10.1038/nrn2762

15. Diekelmann, S., Born, J., & Wagner, U. (2010). Sleep enhances false memories depending on general memory performance. Behavioural Brain Research, 208(2), 425–429.

16. Diering, G. H., & Huganir, R. L. (2018). The AMPA receptor code of synaptic plasticity. Neuron, 100(2), 314–329.

17. Dworak, M., Wiater, A., Alfer, D., Stephan, E., Hollmann, W., & Strüder, H. K. (2008). Increased slow wave sleep and reduced stage 2 sleep in children depending on exercise intensity. Sleep Medicine, 9(3), 266–272. 10.1016/j.sleep.2007.04.017

18. Ellis, B. W., Johns, M. W., Lancaster, R., Raptopoulos, P., Angelopoulos, N., & Priest, R. G. (1981). The St. Mary’s Hospital sleep questionnaire: A study of reliability. Sleep, 4(1), 93–97.

19. Feld, G. B., & Born, J. (2017). Sculpting memory during sleep: Concurrent consolidation and forgetting. Current Opinion in Neurobiology, 44, 20–27. 10.1016/j.conb.2017.02.012

20. Feld, G. B., & Born, J. (2020). Neurochemical mechanisms for memory processing during sleep: Basic findings in humans and neuropsychiatric implications. Neuropsychopharmacology, 45(1), 31–44. 10.1038/s41386-019-0490-9

21. Feld, G. B., & Diekelmann, S. (2020). Building the Bridge: Outlining Steps Toward an Applied Sleep-and-Memory Research Program. Current Directions in Psychological Science, 29(6), 554–562. 10.1177/0963721420964171

22. Feld, G. B., Weis, P. P., & Born, J. (2016). The limited capacity of sleep-dependent memory consolidation. Frontiers in Psychology, 7, 1368.

23. Frimpong, E., Mograss, M., Zvionow, T., & Dang-Vu, T. T. (2021). The effects of evening high-intensity exercise on sleep in healthy adults: A systematic review and meta-analysis. Sleep Medicine Reviews, 60, 101535. 10.1016/j.smrv.2021.101535

24. Frimpong, E., Mograss, M., Zvionow, T., Paez, A., Aubertin-Leheudre, M., Bherer, L., Pepin, V., Robertson, E. M., & Dang-Vu, T. T. (2023). Acute evening high-intensity interval training may attenuate the detrimental effects of sleep restriction on long-term declarative memory. SLEEP, 46(7), zsad119. 10.1093/sleep/zsad119

25. Frisch, N., Heischel, L., Wanner, P., Kern, S., Gürsoy, Ç. N., Roig, M., Feld, G. B., & Steib, S. (2024). An acute bout of high-intensity exercise affects nocturnal sleep and sleep-dependent memory consolidation. Journal of Sleep Research, 33(4), e14126. 10.1111/jsr.14126

26. Frith, E., Sng, E., & Loprinzi, P. D. (2017). Randomized controlled trial evaluating the temporal effects of high-intensity exercise on learning, short-term and long-term memory, and prospective memory. European Journal of Neuroscience, 46(10), 2557–2564. 10.1111/ejn.13719

27. Goodman, J. M., Thomas, S. G., & Burr, J. (2011). Evidence-based risk assessment and recommendations for exercise testing and physical activity clearance in apparently healthy individuals ^1^ This paper is one of a selection of papers published in this Special Issue, entitled Evidence-based risk assessment and recommendations for physical activity clearance, and has undergone the Journal’s usual peer review process. *Applied Physiology*, Nutrition, and Metabolism, 36(S1), S14–S32. 10.1139/h11-048

28. Hayashi, Y., Nishihira, Y., & Higashiura, T. (2014). The effects of different intensities of exercise on night sleep. Advances in Exercise and Sports Physiology, 20(1), 19–24.

29. Hinard, V., Mikhail, C., Pradervand, S., Curie, T., Houtkooper, R. H., Auwerx, J., Franken, P., & Tafti, M. (2012). Key electrophysiological, molecular, and metabolic signatures of sleep and wakefulness revealed in primary cortical cultures. Journal of Neuroscience, 32(36), 12506–12517.

30. Hoddes, E., Zarcone, V., Smythe, H., Phillips, R., & Dement, W. C. (1973). Quantification of Sleepiness: A New Approach. Psychophysiology, 10(4), 431–436. 10.1111/j.1469-8986.1973.tb00801.x

31. Jentsch, V. L., & Wolf, O. T. (2020). Acute physical exercise promotes the consolidation of emotional material. Neurobiology of Learning and Memory, 173, 107252.

32. Jones, A. P., Bryant, N. B., Robert, B. M., Mullins, T. S., Trumbo, M. C., Ketz, N. A., Howard, M. D., Pilly, P. K., & Clark, V. P. (2023). Closed-loop tACS delivered during slow-wave sleep reduces retroactive interference on a paired-associates learning task. Brain Sciences, 13(3), 468.

33. Kao, S.-C., Drollette, E. S., Ritondale, J. P., Khan, N., & Hillman, C. H. (2018). The acute effects of high-intensity interval training and moderate-intensity continuous exercise on declarative memory and inhibitory control. Psychology of Sport and Exercise, 38, 90–99. 10.1016/j.psychsport.2018.05.011

34. Kredlow, M. A., Capozzoli, M. C., Hearon, B. A., Calkins, A. W., & Otto, M. W. (2015). The effects of physical activity on sleep: A meta-analytic review. Journal of Behavioral Medicine, 38(3), 427–449. 10.1007/s10865-015-9617-6

35. Kubitz, K. A., Landers, D. M., Petruzzello, S. J., & Han, M. (1996). The Effects of Acute and Chronic Exercise on Sleep: A Meta-Analytic Review. Sports Medicine, 21(4), 277–291. 10.2165/00007256-199621040-00004

36. Kuznetsova, A., Brockhoff, P. B., & Christensen, R. H. B. (2017). lmerTest package: Tests in linear mixed effects models. Journal of Statistical Software, 82(13). https://orbit.dtu.dk/en/publications/lmertest-package-tests-in-linear-mixed-effects-models

37. Lakens, D. (2022). Sample Size Justification. Collabra: Psychology, 8(1), 33267. 10.1525/collabra.33267

38. Lakens, D. (2024). When and How to Deviate from a Preregistration. Collabra: Psychology, 10(1). https://online.ucpress.edu/collabra/article/10/1/117094/200749

39. Lechner, H. A., Squire, L. R., & Byrne, J. H. (1999). 100 years of consolidation— Remembering Müller and Pilzecker. Learning & Memory, 6(2), 77–87.

40. Lenth, R. (2024). Emmeans: Estimated marginal means, aka least-squares means. R package version 1.*9*. 0. 2023.

41. Liu, Z.-W., Faraguna, U., Cirelli, C., Tononi, G., & Gao, X.-B. (2010). Direct evidence for wake-related increases and sleep-related decreases in synaptic strength in rodent cortex. Journal of Neuroscience, 30(25), 8671–8675.

42. Lopes da Cunha, P., Ramírez Butavand, D., Chisari, L. B., Ballarini, F., & Viola, H. (2018). Exams at classroom have bidirectional effects on the long-term memory of an unrelated graphical task. Npj Science of Learning, 3(1), 19.

43. Loprinzi, P. D., Blough, J., Crawford, L., Ryu, S., Zou, L., & Li, H. (2019). The temporal effects of acute exercise on episodic memory function: Systematic review with meta-analysis. Brain Sciences, 9(4), 87.

44. Mander, B. A., Santhanam, S., Saletin, J. M., & Walker, M. P. (2011). Wake deterioration and sleep restoration of human learning. Current Biology, 21(5), R183–R184. 10.1016/j.cub.2011.01.019

45. Matuschek, H., Kliegl, R., Vasishth, S., Baayen, H., & Bates, D. (2017). Balancing Type I error and power in linear mixed models. Journal of Memory and Language, 94, 305–315.

46. Mograss, M., Crosetta, M., Abi-Jaoude, J., Frolova, E., Robertson, E. M., Pepin, V., & Dang-Vu, T. T. (2020). Exercising before a nap benefits memory better than napping or exercising alone. Sleep, 43(9), zsaa062. 10.1093/sleep/zsaa062

47. Myllymäki, T., Kyröläinen, H., Savolainen, K., Hokka, L., Jakonen, R., Juuti, T., Martinmäki, K., Kaartinen, J., Kinnunen, M.-L., & Rusko, H. (2011). Effects of vigorous late-night exercise on sleep quality and cardiac autonomic activity: Late-night exercise and sleep. Journal of Sleep Research, 20(1pt2), 146–153. 10.1111/j.1365-2869.2010.00874.x

48. Nebe, S., Reutter, M., Baker, D. H., Bölte, J., Domes, G., Gamer, M., Gärtner, A., Gießing, C., Gurr, C., & Hilger, K. (2023). Enhancing precision in human neuroscience. Elife, 12, e85980.

49. Ong, J. L., Lau, T. Y., Lee, X. K., Van Rijn, E., & Chee, M. W. L. (2020). A daytime nap restores hippocampal function and improves declarative learning. Sleep, 43(9), zsaa058. 10.1093/sleep/zsaa058

50. Ostadan, F., Centeno, C., Daloze, J.-F., Frenn, M., Lundbye-Jensen, J., & Roig, M. (2016). Changes in corticospinal excitability during consolidation predict acute exercise-induced off-line gains in procedural memory. Neurobiology of Learning and Memory, 136, 196–203.

51. Park, I., Díaz, J., Matsumoto, S., Iwayama, K., Nabekura, Y., Ogata, H., Kayaba, M., Aoyagi, A., Yajima, K., Satoh, M., Tokuyama, K., & Vogt, K. E. (2021). Exercise improves the quality of slow-wave sleep by increasing slow-wave stability. Scientific Reports, 11(1), 4410.10.1038/s41598-021-83817-6

52. Ramírez Butavand, D., Hirsch, I., Tomaiuolo, M., Moncada, D., Viola, H., & Ballarini, F. (2020). Novelty improves the formation and persistence of memory in a naturalistic school scenario. Frontiers in Psychology, 11, 48.

53. Rasch, B., & Born, J. (2013). About Sleep’s Role in Memory. Physiological Reviews, 93(2), 681–766. 10.1152/physrev.00032.2012

54. Roach, G. D., Dawson, D., & Lamond, N. (2006). Can a Shorter Psychomotor Vigilance Task Be Usedas a Reasonable Substitute for the Ten-Minute Psychomotor Vigilance Task? Chronobiology International, 23(6), 1379–1387. 10.1080/07420520601067931

55. Robertson, E. M. (2012). New insights in human memory interference and consolidation. Current Biology, 22(2), R66–R71.

56. Roig, M., Cristini, J., Parwanta, Z., Ayotte, B., Rodrigues, L., De Las Heras, B., Nepveu, J.-F., Huber, R., Carrier, J., Steib, S., Youngstedt, S. D., & Wright, D. L. (2022). Exercising the Sleepy-ing Brain: Exercise, Sleep, and Sleep Loss on Memory. Exercise and Sport Sciences Reviews, 50(1), 38–48. 10.1249/JES.0000000000000273

57. Roig, M., Nordbrandt, S., Geertsen, S. S., & Nielsen, J. B. (2013). The effects of cardiovascular exercise on human memory: A review with meta-analysis. Neuroscience & Biobehavioral Reviews, 37(8), 1645–1666. 10.1016/j.neubiorev.2013.06.012

58. Roig, M., Skriver, K., Lundbye-Jensen, J., Kiens, B., & Nielsen, J. B. (2012). *A single bout of exercise improves motor memory*. https://journals.plos.org/plosone/article?id=10.1371/journal.pone.0044594

59. Roig-Hierro, E., & Batalla, A. (2023). Acute exercise on complex motor memory retention: The role of task cognitive demands. International Journal of Sport and Exercise Psychology, 1–15. 10.1080/1612197X.2023.2278522

60. Schwenk, J., Baehrens, D., Haupt, A., Bildl, W., Boudkkazi, S., Roeper, J., Fakler, B., & Schulte, U. (2014). Regional diversity and developmental dynamics of the AMPA-receptor proteome in the mammalian brain. Neuron, 84(1), 41–54.

61. Stickgold, R., & Walker, M. P. (2013). Sleep-dependent memory triage: Evolving generalization through selective processing. Nature Neuroscience, 16(2), 139–145. 10.1038/nn.3303

62. Suzuki, A., Yanagisawa, M., & Greene, R. W. (2020). Loss of *Arc* attenuates the behavioral and molecular responses for sleep homeostasis in mice. Proceedings of the National Academy of Sciences, 117(19), 10547–10553. 10.1073/pnas.1906840117

63. Thomas, C., Jones, H., Whitworth-Turner, C., & Louis, J. (2020). High-intensity exercise in the evening does not disrupt sleep in endurance runners. European Journal of Applied Physiology, 120(2), 359–368. 10.1007/s00421-019-04280-w

64. Thomas, R., Johnsen, L. K., Geertsen, S. S., Christiansen, L., Ritz, C., Roig, M., & Lundbye-Jensen, J. (2016). Acute exercise and motor memory consolidation: The role of exercise intensity. PloS One, 11(7), e0159589.

65. Tononi, G., & Cirelli, C. (2003). Sleep and synaptic homeostasis: A hypothesis. Brain Research Bulletin, 62(2), 143–150. 10.1016/j.brainresbull.2003.09.004

66. Tononi, G., & Cirelli, C. (2006). Sleep function and synaptic homeostasis. Sleep Medicine Reviews, 10(1), 49–62. 10.1016/j.smrv.2005.05.002

67. Tononi, G., & Cirelli, C. (2014). Sleep and the Price of Plasticity: From Synaptic and Cellular Homeostasis to Memory Consolidation and Integration. Neuron, 81(1), 12–34. 10.1016/j.neuron.2013.12.025

68. Torsvall, L., Åkerstedt, T., & Göran, L. (1984). Effects on sleep stages and EEG power density of different degrees of exercise in fit subjects. Electroencephalography and Clinical Neurophysiology, 57(4), 347–353.

69. Tucker, M. A., & Fishbein, W. (2008). Enhancement of declarative memory performance following a daytime nap is contingent on strength of initial task acquisition. Sleep, 31(2), 197–203.

70. Van Der Werf, Y. D., Altena, E., Schoonheim, M. M., Sanz-Arigita, E. J., Vis, J. C., De Rijke, W., & Van Someren, E. J. W. (2009). Sleep benefits subsequent hippocampal functioning. Nature Neuroscience, 12(2), 122–123. 10.1038/nn.2253

71. van Dongen, E. V., Kersten, I. H., Wagner, I. C., Morris, R. G., & Fernández, G. (2016). Physical exercise performed four hours after learning improves memory retention and increases hippocampal pattern similarity during retrieval. Current Biology, 26(13), 1722–1727.

72. Vyazovskiy, V. V., Cirelli, C., Pfister-Genskow, M., Faraguna, U., & Tononi, G. (2008). Molecular and electrophysiological evidence for net synaptic potentiation in wake and depression in sleep. Nature Neuroscience, 11(2), 200–208.

73. Wanner, P., Cheng, F.-H., & Steib, S. (2020). Effects of acute cardiovascular exercise on motor memory encoding and consolidation: A systematic review with meta-analysis. Neuroscience & Biobehavioral Reviews, 116, 365–381.

74. Yoo, S.-S., Hu, P. T., Gujar, N., Jolesz, F. A., & Walker, M. P. (2007). A deficit in the ability to form new human memories without sleep. Nature Neuroscience, 10(3), 385–392. 10.1038/nn1851

75. Yoshida, H., Ishikawa, T., Shiraishi, F., & Kobayashi, T. (1998). Effects of the timing of exercise on the night sleep. Psychiatry and Clinical Neurosciences, 52(2), 139–140. 10.1111/j.1440-1819.1998.tb00994.x

76. Youngstedt, S. D., O’Connor, P. J., & Dishman, R. K. (1997). The Effects of Acute Exercise on Sleep: A Quantitative Synthesis. Sleep, 20(3), 203–214. 10.1093/sleep/20.3.203

