## Supplementary Material for "High intensity exercise before sleep boosts memory encoding the next morning"

[1]Human Movement, Training and Active Aging Department, Institute of Sports and Sports Science, Heidelberg University, Heidelberg, Germany.
[2]Clinical Psychology, Central Institute of Mental Health, Medical Faculty Mannheim, Heidelberg University, Mannheim, Germany.
[3]Addiction Behavior and Addiction Medicine, Central Institute of Mental Health, Medical Faculty Mannheim, Heidelberg University, Mannheim, Germany.
[4]Psychiatry and Psychotherapy, Central Institute of Mental Health, Medical Faculty Mannheim, Heidelberg University, Mannheim, Germany.
[5]Psychological Institute, Heidelberg University, Heidelberg, Germany.
[6]German Center for Mental Health (DZPG).
[7]Network Aging Research, Heidelberg University
[†] SS and GF contributed equally to this work
*

### Supplementary analysis 1

The results of the PVT analysis can be found in Table 1. The best-fitting model was: *reaction speed ~ condition + test + (1 + condition + test | subject)*. The reference level for the predictor condition was control and for the predictor test was E1.

| | df | *t*-value | *p*-value |
|---|---|---|---|
| **HIIT** | 36.56 | 0.91 | 0.369 |
| **MICT** | 33.97 | 1.19 | 0.242 |
| **E2** | 73.48 | -3.94 | <0.001 |
| **R24h** | 46.94 | -0.64 | 0.526 |

**Table 1**: Fixed effects derived from the analysis of reaction speed measured in the Psychomotor Vigilance Task (PVT).

### Supplementary analysis 2

The sleep results for the second night (before R24h) can be found in Table 2. The best-fitting model for each sleep variable was: *sleep variable ~ condition + (1 | subject)*.

| | Control | | HIIT | | | MICT | | |
|---|---|---|---|---|---|---|---|---|
| Variable | Mean | SD | Mean | SD | *p*-value | Mean | SD | *p*-value |
| **Latency** (min) | 12.4 | 13.9 | 10.5 | 15.7 | 0.581 | 15.7 | 16.2 | 0.364 |
| **Efficiency** (%) | 79.4 | 8.5 | 78.5 | 9.7 | 0.680 | 77.8 | 7.9 | 0.532 |
| **TTB** (min) | 485 | 66 | 484 | 63 | 0.816 | 487 | 66 | 0.878 |
| **TST** (min) | 385 | 62 | 379 | 64 | 0.570 | 378 | 63 | 0.821 |
| **WASO** (min) | 88.2 | 43.4 | 92.6 | 42.5 | 0.493 | 94.6 | 48.9 | 0.805 |
| **#awakenings** | 28.5 | 12.3 | 31.9 | 9.8 | 0.114 | 31.4 | 10.6 | 0.061 |
| **Avg. Awakenings** (min) | 3.2 | 1.3 | 2.9 | 1.0 | 0.296 | 3.0 | 1.2 | 0.130 |

**Table 2**: **Sleep Variables for the second night (before R24h).** The mean and standard deviation (SD) values for each sleep variable (Latency, Efficiency, Total Time in Bed [TTB], and Total Sleep Time [TST], Wake After Sleep Onset [WASO], number [#] of awakenings, and average awakenings) are shown for each condition. In addition, the *p*-values from the comparisons between conditions are displayed.

### Supplementary analysis 3

We performed the pre-registered correlation between sleep efficiency and memory performance. There was no significant correlation between these two variables ($r$ = -0.002, $p$ = 0.981).

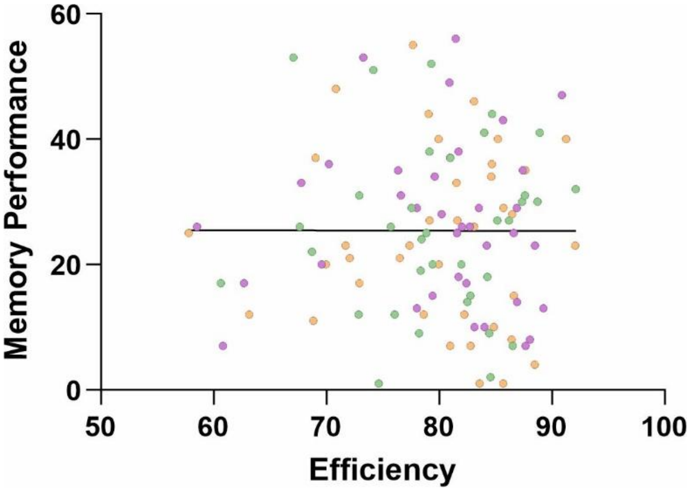


**Figure 1**: Correlation between memory performance and sleep efficiency.

### Supplementary analysis 4

For analyzing the effect of exercise on the different phases of encoding we ran the following best-fitting model: *memory performance (0 or 1) ~ condition * encoding time + (1 + condition + encoding time | subject) + (1 | item).* The results for E1 and R24h tests are reported in the main manuscript. The results of additional pairwise comparisons of the E1 analysis can be found in Table 3.

The analysis for E2 was run with the following model: *memory performance (0 or 1) ~ condition * encoding time + (1 + condition | subject) + (1 | item)* because the model used for the other two analyses presented boundary singular fit in the E2 data. The results for E2 can be found in Table 4.

| Contrast | Estimate | SE | *z* - ratio | *p* - value |
|---|---|---|---|---|
| Control – HIIT (early) | -0.46 | 0.14 | -3.35 | 0.011* |
| Control – MICT (early) | -0.26 | 0.17 | -1.57 | 0.621 |
| HIIT – MICT (early) | 0.20 | 0.14 | 1.50 | 0.666 |
| Control – HIIT (late) | -0.07 | 0.14 | -0.50 | 0.996 |
| Control – MICT (late) | 0.16 | 0.17 | 0.95 | 0.933 |
| HIIT – MICT (late) | 0.23 | 0.14 | 1.64 | 0.569 |

**Table 3**: Pairwise comparisons results from the estimated marginal means for the analysis of the effect of exercise on the different phases of encoding (early and late) on E1 data.

| | β | SE | *z* | *p* - value |
|---|---|---|---|---|
| Condition (Control vs. HIIT) : encoding time (early vs. late) | -0.25 | 0.13 | -1.91 | 0.057. |
| Condition (Control vs. MICT) : encoding time (early vs. late) | -0.20 | 0.13 | -1.50 | 0.132 |
| Condition: HIIT | 0.35 | 0.15 | 2.34 | 0.019* |
| Condition: MICT | 0.11 | 0.16 | 0.69 | 0.490 |
| Encoding time: late | 0.05 | 0.09 | 0.61 | 0.545 |

**Table 4**: Statistical parameters of the analysis including early and late encoding in E2 tests. The reference level for the predictor condition was control and for the predictor encoding_time was early.

### Supplementary analysis 5

For analyzing the effect of exercise on low- and high-performing participants we ran the following best-fitting model: *memory performance (0 or 1) ~ condition * performer + (1 + condition | subject) + (1 | item).* The results for E1 test are reported in the main manuscript.
The results of additional pairwise comparisons of the E1 analysis can be found in Table 5.
The analysis for E2 can be found in Table 6 and for R24h in Table 7.

| **Contrast** | **Estimate** | **SE** | ***z* - ratio** | ***p* - value** |
|---|---|---|---|---|
| Control – HIIT (low) | -0.52 | 0.16 | -3.15 | 0.020* |
| Control – MICT (low) | -0.25 | 0.22 | -1.15 | 0.859 |
| HIIT – MICT (low) | 0.27 | 0.17 | 1.56 | 0.628 |
| Control – HIIT (high) | -0.01 | 0.16 | -0.05 | 1.000 |
| Control – MICT (high) | 0.12 | 0.20 | 0.60 | 0.991 |
| HIIT – MICT (high) | 0.13 | 0.16 | 0.78 | 0.971 |

**Table 5**: Pairwise comparisons results from the estimated marginal means for the analysis of the effect of exercise on low- and high-performing participants on E1 data.

| | **β** | **SE** | ***z*** | ***p* - value** |
|---|---|---|---|---|
| Condition (Control vs. HIIT) : performer (low vs. high) | -0.37 | 0.27 | -1.41 | 0.160 |
| Condition (Control vs. MICT) : performer (low vs. high) | -0.37 | 0.29 | -1.26 | 0.208 |
| Condition: HIIT | 0.42 | 0.18 | 2.35 | 0.019* |
| Condition: MICT | 0.20 | 0.21 | 0.96 | 0.338 |
| Performer: high | 1.95 | 0.29 | 6.82 | <0.001*** |

**Table 6**: Statistical parameters of the analysis including low- and high-performing participants in E2 tests. The reference level for the predictor condition was control and for the predictor performer was low.

| | **β** | **SE** | ***z*** | ***p* - value** |
|---|---|---|---|---|
| Condition (Control vs. HIIT) : performer (low vs. high) | -0.35 | 0.27 | -1.26 | 0.208 |
| Condition (Control vs. MICT) : performer (low vs. high) | -0.09 | 0.33 | -0.28 | 0.782 |
| Condition: HIIT | 0.48 | 0.19 | 2.47 | 0.014* |
| Condition: MICT | 0.15 | 0.24 | 0.62 | 0.535 |
| Performer: high | 1.88 | 0.27 | 6.96 | <0.001*** |

**Table 7**: Statistical parameters of the analysis including low- and high-performing participants in R24h tests. The reference level for the predictor condition was control and for the predictor performer was low. Related to Fig. 4.
